# The emerging threat of hot drought in Western Australia

**DOI:** 10.1101/2025.01.22.633981

**Authors:** Stanley Mastrantonis, Amanda R Bourne

## Abstract

Droughts severely affect our environments, biodiversity, production systems and communities. When assessing droughts, we generally consider deficits in rainfall, subsurface water, water for crops and water for societies, economies and the environment. Due to climate change, increasing temperatures and more frequent and prolonged drought events are occurring in Australia and globally. When droughts and extreme temperatures occur together in the same space and time, it results in hot droughts, a phenomenon that can be devastating for flora, fauna and society. Here, we map hot drought events across Western Australia since 1889. Our results indicate that significant hot droughts have become more frequent, with the most severe hot droughts on record occurring within the last five years. Additionally, hot droughts within the last four decades, and particularly within the last two decades, affected a significantly larger area than the average for the historical record. With 2024 already on track to be the driest and hottest year on record in southwest Western Australia, natural resource managers must prepare for the increasing frequency and severity of hot droughts.

**Implication for Managers:**

- Australia and the world are experiencing more frequent and intense droughts and heatwaves. Hot droughts, characterised by simultaneous extreme water scarcity and extreme high temperatures, are particularly devastating.
- We mapped hot drought events in Western Australia since 1889, revealing a recent surge in occurrence concentrated in the most recent five years. Statistical testing indicates that hot droughts within the most recent 20 and 40 years were significantly larger in extent compared to the historical average.
- With the first half of 2024 exceptionally dry and hot, proactive strategies are imperative for natural resource managers.

## 1. Introduction

Droughts are among the costliest natural disasters, with changes in hydrological regimes having far-reaching impacts, including threatening life and food security and increasing bushfire risk (Wasko et al. 2021). Drought’s economic and health impacts are particularly prevalent in Australia, ranking 5^th^ for economic losses and 15^th^ for the number of people affected (Kirono et al. 2020). Due to warming climates, drought trends vary spatially, with the tropical regions of Australia and the world experiencing wetter climates while temperate regions are experiencing rainfall declines(Wasko et al. 2021).

The Mediterranean climate region of the South West of Western Australia (SWWA) is a global biodiversity hotspot that supports important agriculture and most of the state’s population (Myers et al. 2000; Wang et al. 2018). The SWWA has experienced hotter and drier climates, particularly over the last 40 years (Wasko et al. 2021; Mcfarlane et al. 2020; Fletcher et al. 2020). Rainfall in the cool season (May-October) contributes to 70% of the total annual rainfall for the SWWA and rain follows cold fronts from northwards movements of the subtropical ridge (Timbal and Drosdowsky 2013). Warm season rainfall is linked to thunderstorms and cyclones moving down from the tropics (Rauniyar et al. 2023).

Prior to 1960, cool season rainfall exhibited high interannual variability with extreme fluctuations between very wet and very dry cool seasons. After the 1960s the frequency of very wet seasons started to decline, and this drying trend has become more pronounced since the 1990s (Rauniyar et al. 2023) temperatures have increased by 0.5°C since the 1960s for the SWWA, while evaporation has increased by 200 mm per annum and both streamflows and surface water runoff have also declined (Mcfarlane et al. 2020). These changes in climate and hydrological regimes were originally attributed to natural cycles but as water availability became increasingly limited and trends became entrenched, anthropogenic climate change was recognised as the main driver (Bates et al. 2010). Factors such as significant increases in greenhouse gasses, changes in atmospheric aerosols, land transformation and human development contribute to increasingly limited water availability in the SWWA, with more severe and frequent droughts expected in the future (Hansen et al. 2023; Elkouk et al. 2022; Vicente-Serrano et al. 2015; Mcfarlane et al. 2020; Bates et al. 2010). In addition to the anthropogenic factors influencing climate, rainfall in southern Australia is strongly influenced by variations in large-scale atmospheric circulation such as the Southern Annular Mode (SAM), the Indian Ocean Dipole (IOD) and the El Niño-Southern Oscillation (ENSO). Rainfall declines in SWWA have been attributed, in part, to a positive trend of the SAM, where strong westerly winds shift polewards and reduce cold fronts during the cool season (Hope et al. 2015). Large-scale Land Cover Change (LCC) has also been attributed to localised warming and drying in the SWWA, and the interaction between large-scale LCC and increasing greenhouse gas emissions has led to significant declines in water availability for the SWWA (Pitman et al. 2004).

Though anthropogenic climate change is one driving factor for declining rainfall in the SWWA, external forcings also contributed to this recent drying trend (Rauniyar et al. 2023). Future climate outlooks from ensemble circulation models such as the Coupled Model Intercomparison Project (CMIP) all forecast increased drying and warming for sub-tropical regions due to increasing greenhouse gas emissions. For the SWWA, winter and spring rainfalls are expected to decrease by 20% to 60% by the end of the century under the Representative Concentration Pathway 8.5 (RCP 8.5) scenario, leading to more prolonged, frequent and severe droughts for the region (Hope et al. 2015).

Like droughts, heat waves can be defined as prolonged periods of unusually high heat stress (Robinson 2001). They can have severe effects on societies, production systems and natural environments, including coastal and marine environments (Robinson 2001; Wreford and Adger 2010; Oliver et al. 2018). In Australia, heatwaves have killed more people than all other natural hazards combined (Xiao et al. 2017). Since 2000, the frequency, duration and severity of heatwaves have significantly increased in Australia, particularly in recent decades, resulting in vegetation die-offs, population declines for native fauna, crop failures and health impacts on human societies (Bourne et al. 2023, 2020; Mastrantonis et al. 2019; Zhang, Chen, and Chen 2023; Allen, Breshears, and McDowell 2015; Hartmann et al. 2022, 2018; Jyoteeshkumar reddy, Perkins-Kirkpatrick, and Sharples 2021). Land affected by heatwaves is expected to quadruple by 2040, while marine heatwaves now occur 4-5 times more often than historical rates (Ruthrof et al. 2018). A 2011 marine and land heatwave event that occurred across the ocean and coast of SWAA triggered abrupt, synchronous, and multi-trophic ecological disruptions for both marine and terrestrial taxa (Ruthrof et al. 2018).

Similar to droughts, global climate change, resulting from increasing greenhouse gas emissions and human development, is the main factor driving the intensification of heatwave frequency and duration (Hirsch et al. 2019; Trancoso et al. 2020). Natural climate cycles such as the ENSO and IOD can exacerbate heatwave events (Trancoso et al. 2020), and interactions between anthropogenic-induced warming and natural climate cycles likely compound heatwave impacts within years.

Severe droughts and heatwaves have had documented negative effects on many species (Cruz-McDonnell and Wolf 2016; Klockmann and Fischer 2017; Brivio et al. 2019; Mastrantonis et al. 2019; Anderegg et al. 2019; Sharpe et al. 2019) and production systems (Lawes and Kingwell 2012; Souther et al. 2019; Jiao et al. 2020; Coughlan De Perez et al. 2023), including mass mortality events, population decline and crop failure (Read 2004; McKechnie and Wolf 2010; Saunders et al. 2011; Allen et al. 2015; Ratnayake et al. 2019; Brás et al. 2021; Pattinson et al. 2022). Hot droughts are emerging compound events in which extremely high temperatures co-occur with extremely low rainfall, with the potential to exacerbate the impacts of both types of extreme events on natural ecosystems and agricultural production systems alike in the context of global climate change (Albright et al. 2010; Overpeck 2013; Conrey et al. 2016; Bourne et al. 2020b; Cohen et al. 2021; Dezetter et al. 2022). Hot droughts are currently breaking records around the world, with many regions experiencing unprecedented and unseasonal high temperatures and low rainfall simultaneously (Le Nohaïc et al. 2017; Kreibich et al. 2022; Lemus-Canovas et al. 2024). As the extent and duration of droughts and heatwaves are expected to increase (Overpeck 2013), the spatiotemporal overlap of these climate extremes will also increase. Thus, the severity, extent and duration of hot droughts will follow suit. Here, we consider historical data on rainfall and temperature across Western Australia from 1889 to 2024 and identify trends in the cooccurrence of extremely dry and extremely hot weather across time and space. We map the spatial extent and frequency of hot droughts in Western Australia over this 135-year period. Finally, we statistically test whether recent incidences of hot drought differ significantly from the historical mean.

## 2. Methods

Drought can simply be defined as a “lack of rain” (Hughes et al. 2022). But context matters, and drought can affect agriculture, hydrological cycles, ecological communities and societies. Thus, drought could be better defined as “a prolonged period of abnormally dry conditions when the amount of water available is insufficient to meet the usual needs of local socio-ecological systems” (Bourne et al. 2023). Nevertheless, drought is generally contextualised based on water deficits, and monitoring the occurrence of drought through derivatives or proxies of water availability, such as rainfall amounts or remotely sensed indices, is common practice (Beguería et al. 2014). In Australia, Gibbs & Maher (1967) established rainfall percentiles as the de-facto metric for monitoring meteorological drought, with state and federal governments adopting the use of rainfall percentiles as a trigger for drought support programs due to their ease of use and the ready availability of data (Hughes et al. 2022).

We collated a time series of gridded climate data sourced through SILO Longpaddock (Jeffrey et al. 2001). SILO is an initiative by the Queensland state government to provide up-to-date point and interpolated Australian climate data for various measurements across the continent from 1889 onwards. Specifically, we downloaded historical maximum temperatures (°C) and monthly rainfall (mm) from SILOs Amazon Web Server buckets from the start of 1889 until May 2024 and subset these data for the extent of Western Australia. We constructed a time series for both variables, computing the 10^th^ percentile for total annual rainfall in WA and the 90^th^ percentile for mean maximum temperature. We compared the climate data with the relevant climate percentile each year. For rainfall, locations in a given year with rainfall below the 10^th^ percentile were classified as in drought for that year. Similarly, locations and years where temperatures exceeded the 90^th^ percentile were classified as heatwaves for that year. Where both drought and heatwaves overlapped in space for a given year, the regions were classified as in hot drought. We considered the area (km^2^) affected by hot drought in each year. Additionally, we aggregated the hot drought time series into 20 and 40-year periods to assess how the extent and frequency of hot drought has changed over time. We used a linear model to test if the extent of recent periods of hot drought (1984-2003 and 2004-2024) differed significantly from the historical mean. Additionally, because the SWWA has experienced significant and accelerating drying over the last 40 years as a result of anthropogenic climate change (Dey et al. 2019; Scanlon and Doncon 2020), we compared the frequency of occurrence of hot droughts in the last 20 years with the 20 years prior.

## 3. Results

For the past century, hot droughts have occurred periodically over relatively small areas (mean area in hot drought per year 1889-2018 = 39,646 km^2^). The years of 2019 and the first half of 2024 present notable exceptions in which very large areas were affected (Figs. 1–2). The extremely high temperatures and abnormally low rainfall in 2019 resulted in a hot drought across most of the state of Western Australia, affecting 1,770,608 km^2^ throughout the arid interior of WA (Fig. 2). The 2019 event was three times larger in extent than the next most extensive hot drought in the preceding century, which affected 517,318 km^2^ in 1944 (Fig. 1). At the time of writing (May 2024), Western Australia was in the midst of a second intense and extensive hot drought, affecting almost the entire coastline as well as large inland areas in the southwest, midwest and north of the state and covered 973,104 km^2^ (Fig. 2). Our analysis showed the area affected by hot drought between 2000 and 2024 was an estimated 120,355 km^2^ larger in extent than the historical average (1889-2000), and this difference was statistically significant (*P =* 0.00386). When comparing hot drought from 1980-2024, the area affected by hot drought was an estimated 70,383km^2^ larger in extent than the historical average, and this difference was significant (*P* = 0.0417).

**Figure 1:**
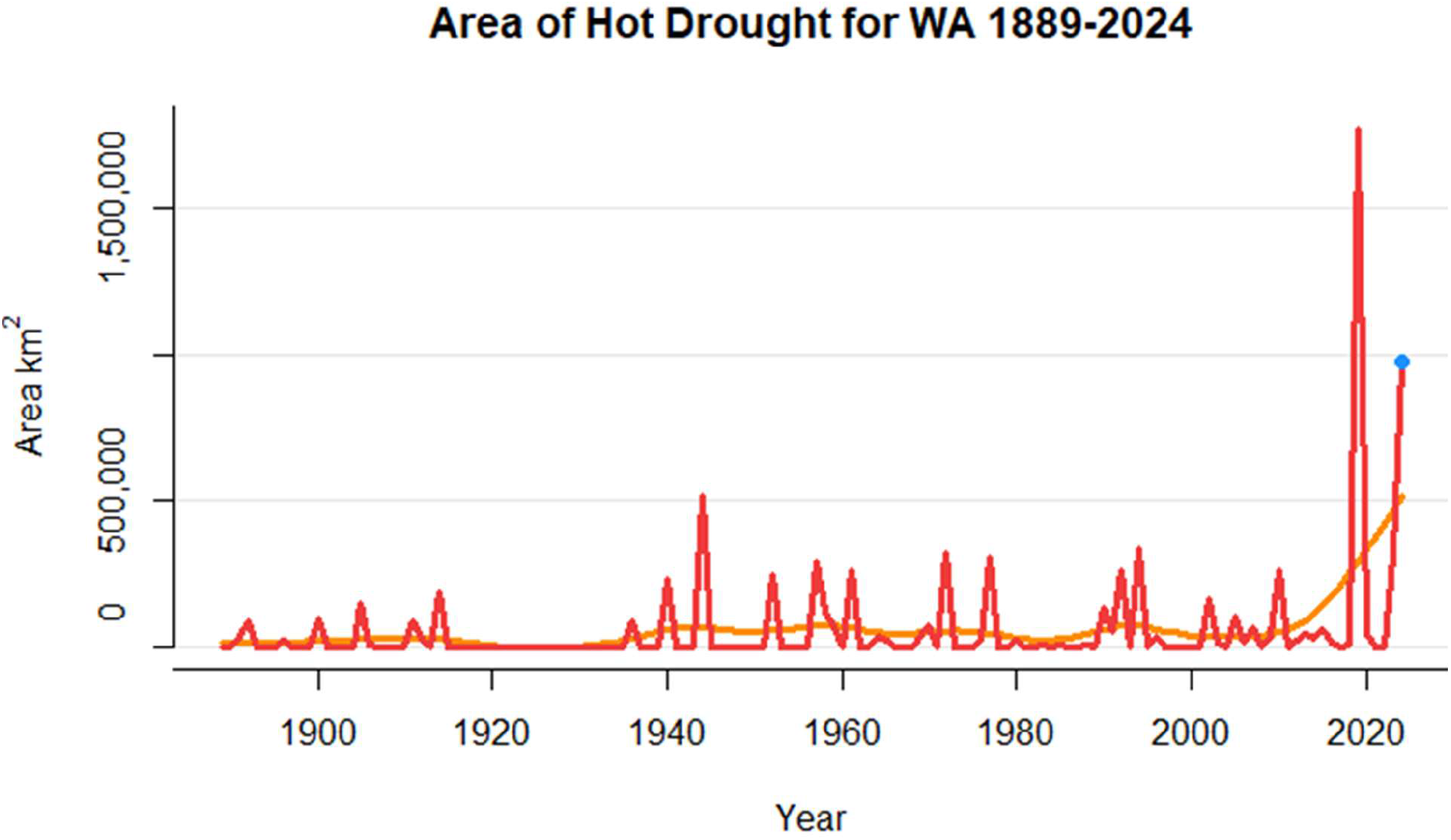
The area (km^2^) of hot drought in WA for the study period. The red lines represent Actual hot drought areas, while the orange hashed line depicts a five-year running average. The blue point is 2024 and shows climate data up until the end of June, the time of writing.

**Figure 2:**
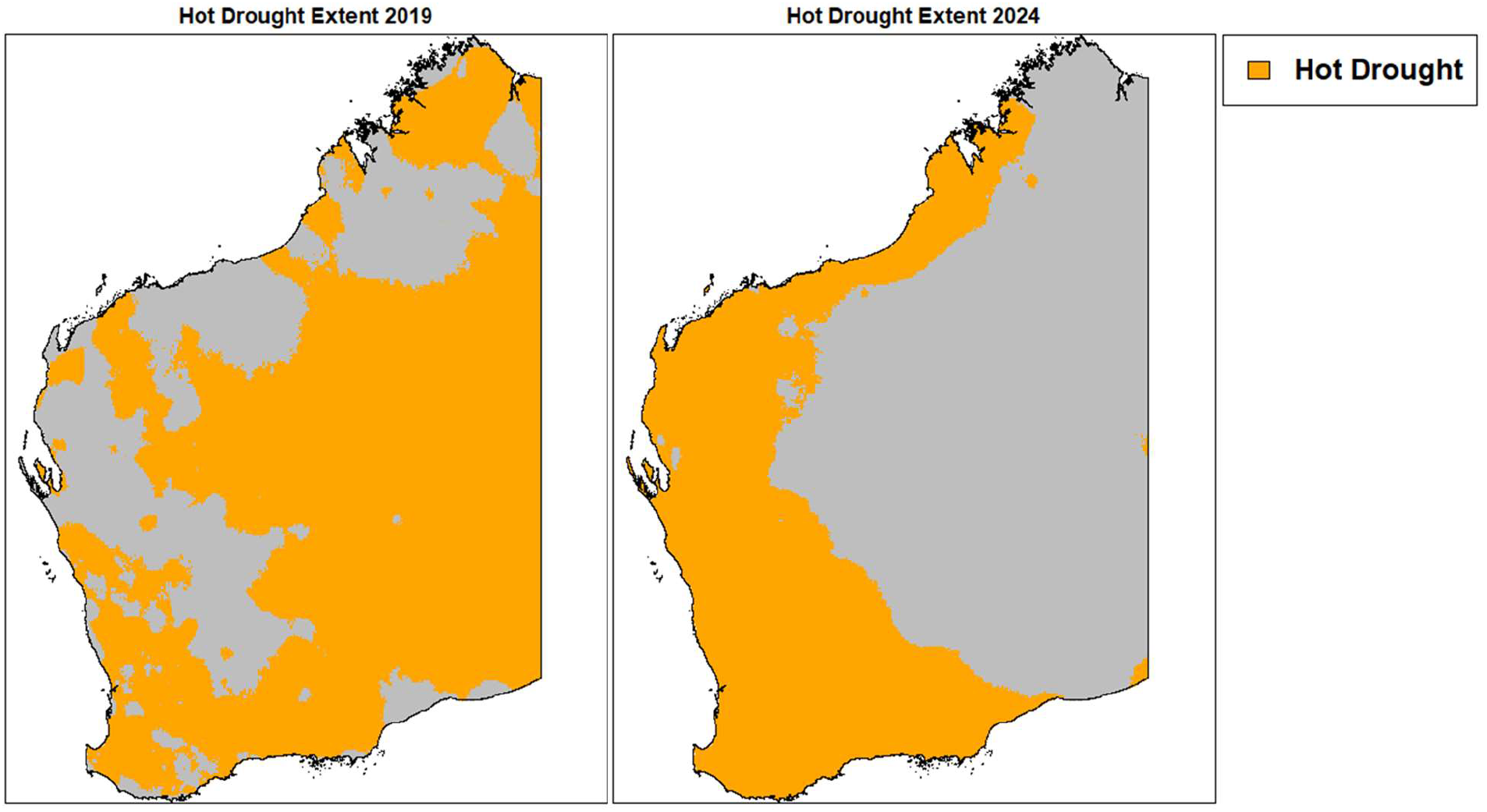
Hot drought extents in Western Australia for 2019 and 2024. Climate data for 2024 extends until the end of May, the time of writing.

## 4. Discussion

Droughts have increased in frequency, intensity and duration in Australia over the last 40 years (Wasko et al. 2021). Simultaneously, heatwaves have increased in frequency, intensity and duration in Australia over the last 40 years (Trancoso et al. 2020). Thus, as the frequency and extent of both drought and heatwaves have increased over the last 40 years, so has the spatiotemporal overlap of these hazards, resulting in increased frequency of hot droughts. Moreover, as unprecedented rates of heatwaves and droughts have occurred in Australia since 2000, more hot droughts have occurred in the last 24 years than in any other period for the span of data (Fig. 1-3). Indeed, in the last two decades, hot droughts have occurred in significantly higher frequencies and over larger areas than they did historically (Fig. 1) and primarily impacted SWWA (Fig. 3). This is an agriculturally important region where the vast majority of the WA population resides (Wang et al. 2018; Fletcher et al. 2020), a designated global biodiversity hotspot (Myers et al. 2000), and an area supporting remnant populations of many endemic threatened plants and animals (Gibson et al. 2004; Woinarski, Burbidge, and) 2014).

**Figure 3:**
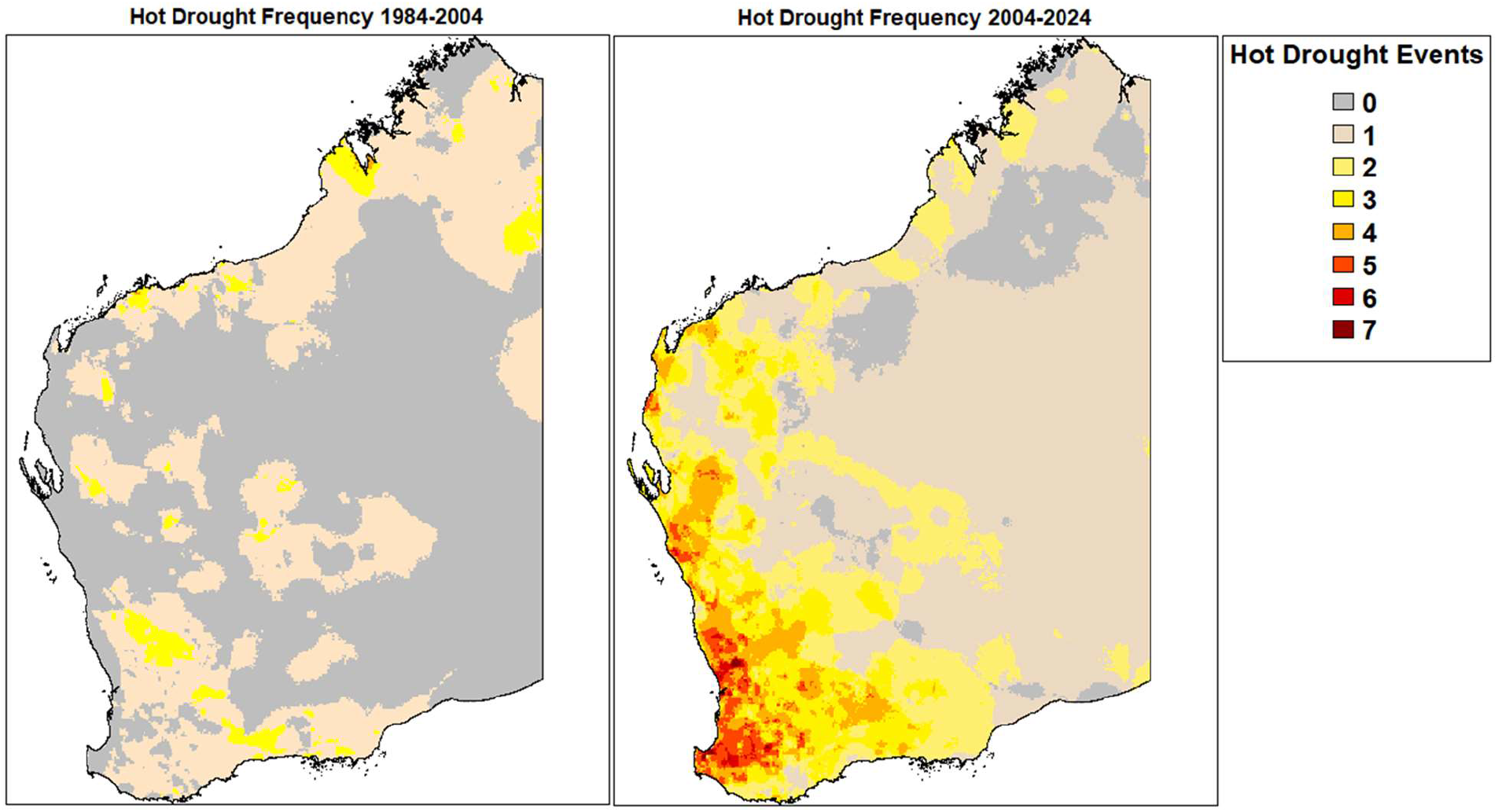
The frequency of hot drought in WA between 1984 and 2004 (left panel) and between 2004 and 2024 (to May; right panel).

Increased greenhouse gasses and external climate forcings have contributed to increasing global mean temperatures (Santer et al. 2013), changes in atmospheric aerosols have shifted Earth’s energy feedbacks (Hansen et al. 2023), and extensive land transformation has altered hydrological and climate regimes around the world (Elkouk et al. 2022; Hirsch et al. 2019; Pitman et al. 2004). These factors, compounding with natural climate cycles such as ENSO, the positive trends of the SAM, and external climate forcings, have all led to the observed increase in frequency and extent of hot drought in Western Australia reported here. Hot droughts have become more frequent and extensive in Western Australia since 2000, and this trend will likely worsen in the future.

The most extreme hot drought in WA to date occurred in 2019, and conditions in the first half of 2024 were also incredibly severe. The frequency of hot droughts has been significantly higher in the most recent two decades than at any other time in the last century. Not only is the frequency and severity of recent hot droughts unprecedented, but the worst affected area (SWWA) coincides with the most biodiverse, populated and agriculturally productive region of the state. The state has already experienced a drying climate in recent decades (Dey et al. 2019; Scanlon and Doncon 2020), and farmers and residents have managed to adapt to increasingly severe droughts (Sprigg et al. 2014; Bourne et al. 2023). However, summer rainfall amounts in 2024 achieved consecutive record lows while temperatures simultaneously and repeatedly achieved record highs. The hot drought over the first half of 2024 has already impacted native vegetation across the state (Matusick et al. 2024), where even urban vegetation is experiencing unprecedented die-offs, and resulted in large declines in several species of reintroduced and extent native fauna (Anonymous, pers comm.). Just as coordinated and dedicated research and effort into marine heat waves occurred in response to a record-breaking event off the coast of Western Australia in 2011 (Holbrook et al. 2020), monitoring, managing and adapting to hot droughts should be prioritised in light of the increasing frequency, severity and spatial extent of such events.

## Notes

https://github.com/StanleyMastrantonis/WA-Drought

### Competing Interest Statement

The authors have declared no competing interest.

